# *Wolbachia w*AlbB inhibit dengue and Zika infection in the mosquito *Aedes aegypti* with an Australian background

**DOI:** 10.1101/2022.03.22.485408

**Authors:** Leon E. Hugo, Gordana Rašić, Andrew J. Maynard, Luke Ambrose, Catherine Liddington, Callum J. E. Thomas, Nisa S. Nath, Melissa Graham, Clay Winterford, B. M. C. Randika Wimalasiri-Yapa, Zhiyong Xi, Nigel W. Beebe, Gregor J. Devine

**Affiliations:** QIMR Berghofer, Brisbane, Queensland, Australia; School of Biological Sciences, University of Queensland, Brisbane, Australia; CSIRO, Brisbane, Queensland, Australia; Department of Medical Laboratory Sciences, The Open University of Sri Lanka, Colombo, Sri Lanka; Department of Microbiology and Molecular Genetics, Michigan State University, East Lansing, MI, USA

**Author notes:** **CORRESPONDING AUTHORS** Leon Hugo, Gregor Devine, QIMR Berghofer Medical Research Institute, 300 Herston Road, Herston Qld 4006 Australia.

**Keywords:** *Wolbachia*, arbovirus, dengue, Zika, *Aedes aegypti*, biological control

## Abstract

Biological control of mosquito vectors using the insect-specific bacteria *Wolbachia* is an emerging strategy for the management of human arboviral diseases. We recently described the development of a strain of *Ae. aegypti* infected with the *Wolbachia* strain *w*AlbB (referred to as the *w*AlbB2-F4 strain) through simple backcrossing of wild type Australian mosquitoes with a *w*AlbB infected *Ae. aegypti* strain from the USA. Field releases of male *w*AlbB2-F4 mosquitoes resulted in the successful suppression of a wild population of mosquitoes in the trial sites by exploiting the strains *Wolbachia-*induced cytoplasmic incompatibility. We now demonstrate that the strain is resistant to infection by dengue and Zika viruses and is genetically similar to endemic Queensland populations. There was a fourfold reduction in the proportion of *w*AlbB2-F4 mosquitoes that became infected following a blood meal containing dengue 2 virus (16.7%) compared to wild type mosquitoes (69.2%) and a 6-7 fold reduction in the proportion of *w*AlbB2-F4 mosquitoes producing virus in saliva following a blood meal containing an epidemic strain of Zika virus (8.7% in comparison to 58.3% in wild type mosquitoes). Restriction-site Associated DNA (RAD) sequencing revealed that *w*AlbB2-F4 mosquitoes have > 98% Australian ancestry, confirming the successful introduction of the *w*AlbB2 infection into the Australian genomic background through backcrossing. Genotypic and phenotypic analyses showed the *w*AlbB2-F4 strain retains the insecticide susceptibility phenotype and genotype of the Australian mosquitoes. We demonstrate that the *Wolbachia w*AlbB2-F4, in addition to being suitable for suppression programs, can be effective in population replacement programs given its high inhibition of virus infection in mosquitoes. The ease at which a target mosquito population can be transfected with *w*AlbB2, while retaining genotypes and phenotypes of the target population, shows the robustness of this strain as a biocontrol agent against the *Ae. aegypti* mosquito itself as well as the pathogens it transmits.

**IMPORTANCE:** Epidemics of arthopod-borne virus (arbovirus) diseases affect millions of people and are becoming more frequent and widespread. A successful strategy to control these diseases is by infecting mosquito populations with benign, insect-specific *Wolbachia* bacteria that render mosquitoes refractory to infection with pathogenic arboviruses. Here we show that a strain of the major mosquito vector *Ae. aegypti* that was infected with *Wolbachia* following a simple back-cross mating procedure is refractory to infection with dengue and Zika viruses. Importantly, the genetic background of the strain is equivalent to the target population, which is important for persistence of the strain and regulatory approval.

---

Arthropod-borne viruses (arboviruses) transmitted by mosquitoes are responsible for global epidemics that are increasing in frequency and geographic scale [1]. Dengue is the most prevalent arboviral disease with 5.2 million cases reported to the WHO in 2019 [2]. The last three decades saw the re-emergence of West Nile, Zika and chikungunya virus diseases in widespread outbreaks [3]. The majority of arboviruses lack vaccines with the exception of Japanese Encephalitis and Yellow Fever viruses. The only commercially available dengue vaccine, Dengvaxia, is only indicated for children nine years and older in populations with a dengue seroprevalence of 70% or greater [4]. In most instances, mosquito management tools are the only options available for combating arbovirus transmission. In many regions, conventional insecticide-based campaigns are compromised by issues of coverage and insecticide resistance and cost concerns in developing countries, so alterative control tools are desperately required. In the last decade, the bacteria *Wolbachia pipientis* has emerged as a major tool for the management of mosquito populations [5].

*Wolbachia* are obligate intracellular endosymbiotic bacteria belonging to the order *Rickettsiales* that are widespread across arthropods thanks to their ability to manipulate the reproductive biology of their hosts [6]. *Wolbachia* infections are transmitted vertically, from female insects to their offspring. *Wolbachia* also induce cytoplasmic incompatibility (CI), an effect where unviable offspring are produced in matings between *Wolbachia*-carrying males and females without *Wolbachia* or with an incompatible *Wolbachia* strain. Thanks to maternal transmission and CI, *Wolbachia* have the ability to invade host populations. *Wolbachia* can also influence the fitness of the host insect, inducing phenotypes that can be either beneficial (nutritional mutualism [7] and pathogen resistance [8, 9]) or deleterious (e.g life shortening[10]). Stable, heritable *Wolbachia* infections in mosquitoes can result in phenotypes that have utility for mosquito population control.

The primary mosquito vector of dengue, Zika and yellow fever viruses, *Aedes aegypti*, is not naturally infected with *Wolbachia*. Stable and heritable *Wolbachia* infections were initially established in *Ae. aegypti* through the microinjection of mosquito eggs [11-13] with *Wolbachia* derived from *Aedes albopictus* (*w*AlbA and *w*AlbB strains) [11], then with bacteria derived from *Drosophila melanogaster* (*w*Mel and *w*MelPop) [12, 13] and *Drosophila simulans* (*w*Au) [14]. The establishment of *Wolbachia* infected strains of *Ae. aegypti* have created new opportunities for mosquito and disease control. Mass releases of *Wolbachia* infected male mosquitoes are being evaluated throughout the globe in a strategy called “population suppression”. At higher densities to the local males, infected males overwhelm the wild population and are responsible for most mating events. As a result of CI, the offspring that result are not viable and die as embryos, causing a subsequent “crash” in *Ae. aegypti* abundance [15-18].

*Wolbachia* infection in some mosquito species may also reduce the capacity of insects to transmit medically important arboviruses and parasites [19, 20]. Induction of pathogen interference by *Wolbachia* in *Ae. aegypti* was first identified from the *w*MelPop strain, which induced strong resistance to dengue and chikungunya viruses and avian malaria parasites [19]. However, sustainable establishment of *w*MelPop at field sites in Vietnam and Australia could not be achieved because of fitness costs that included a substantial reduction to mosquito lifespan [21]. Highly efficient virus transmission blocking is also induced by infections with the more benign *Wolbachia* strains *w*Mel [13], *w*Au [14] and *w*AlbB [22]. By releasing *Wolbachia-*infected male and female mosquitoes in sufficient numbers, a combination of maternal inheritance and CI drives the infection to spread slowly but irreversibly through a mosquito population, resulting in the replacement of the wild type with a more benign and disease refractory form [13, 22-24]. This has been referred to as “population replacement” and various trials around the globe are evaluating the efficacy of population replacement for arbovirus disease control using entomological and epidemiological end points. Deployments of mosquitoes infected with *Wolbachia w*Mel by the World Mosquito Program (formerly Eliminate Dengue) to establish the strain in local mosquito populations have led to significant reductions in dengue incidence in targeted regions, including a 96% reduction in local dengue transmission in northern Australia [25, 26], a 73% reduction in dengue incidence in a quasi-experimental trial in a region of Yogyakarta, Indonesia [27] and 77.1% reduction in dengue incidence across Yogyakarta in a cluster randomised trial [28]. A quasi experimental trial testing the effect of *Wolbachia w*Mel deployment in Niterói, Brazil, observed a 69% reduction in dengue incidence as well as a 56% reduction in chikungunya incidence and a 37% reduction in Zika incidence [29].

The success of *Wolbachia* based population replacement relies on the persistence of the *Wolbachia* infection within local mosquito populations. Data collected over more than a decade since the first deployments of the *Wolbachia w*Mel strain has demonstrated strong persistence of the strain in most regions in which it has been released. *Wolbachia w*Mel has been detected in high prevalence for over eight years in north Queensland [25] and two years in Yogyakarta [27]. *Wolbachia w*Mel prevalences of between 30 and 70% across neighbourhoods of Rio De Janeiro, Brazil, have been achieved over a 131 week period of deployment and monitoring [30]. However, substantial decreases in *Wolbachia w*Mel infection prevalence were recently reported in a local population of *Ae. aegypti* in tropical central Vietnam [31]. By four years after their release, *Wolbachia w*Mel remained in fewer than 5.1% of mosquitoes [31]. The most rapid declines in *Wolbachia* infection prevalence correlated with the onset of the hot dry-season that experiences the hottest average and maximum weekly temperatures. High ambient temperatures typically associated with summer ‘heatwaves’ caused reductions in the density of *w*Mel *Wolbachia* infections in mosquitoes in the laboratory [32, 33] and reductions in prevalence and density following a north Queensland heat wave [15, 34]. *Wolbachia* density is a critical factor defining CI [35], maternal inheritance and pathogen interference [36]. The susceptibility of the *w*Mel strain to heat stress has accelerated the search for alternative *Wolbachia* strains that demonstrate greater tolerance to heat while maintaining the pathogen blocking phenotype.

The *Wolbachia* strain *w*AlbB from the mosquito *Aedes albopictus* was the first to be stably transinfected into *Ae. aegypti*. The strain induced 100% maternal inheritance and strong CI in the new host [11]. In laboratory dengue challenge assays, mosquitoes carrying *w*AlbB were infected with dengue at a significantly lower rate, the titres of dengue virus in infected bodies were reduced and the proportion of mosquitoes with dengue virus in saliva (those potentially capable of transmission) was reduced by at least 37.5% [37]. The *w*AlbB strain reduced dengue transmission potential of mosquitoes to a greater extent than *w*Mel in a robust side-by-side comparison [38]. *Wolbachia w*AlbB is relatively more robust to heat stress than the *w*Mel strain [33]. The *w*AlbB strain was released in Selangor, Malaysia, in 2017 for population replacement in high dengue transmission regions and became established at six trial locations [39]. The strain maintained high infection densities and strong dengue inhibition by 20 months following establishment [22]. The *w*AlbB strain has clear utility for population replacement, particularly in tropical climates.

We recently described the successful suppression of a population of *Ae. aegypti* in northern Australia following mass releases of male mosquitoes from an Australian strain of *Ae aegypti* infected with *Wolbachia w*AlbB [15]. This mosquito strain, referred to as the *Ae. aegypti* wAlbB2-F4 strain, was established by a four-generational backcross whereby *Wolbachia*-free mosquitoes from northern Australia were mated to females from a *w*AlbB infected *Ae. aegypti* strain (WB2). The WB2 strain was originally derived by transinfected *Ae. aegypti* from the USA *Ae. albopictus* B lineage *Wolbachia* (Zhiyong Xi, unpublished). The newly generated *w*AlbB2-F4 displayed complete maternal inheritance of *Wolbachia* and cytoplasmic incompatibility with wild type mosquitoes from Queensland Australia and mosquitoes infected with the *w*Mel strain of *Wolbachia*. Mass releases of male *w*AlbB2-F4 led to overflooding ratios of over 10 infected males per wild male and population suppression of >80% [15]. Here we demonstrate that the *w*AlbB2-F4 strain is also suitable for mosquito population replacement strategies aiming to reduce flavivirus transmission. Females demonstrated strong resistance to infection with dengue 2 virus (DENV-2) and Zika virus (ZIKV) and are equivalent to north Queensland mosquitoes genotypically. This genetic similarity was required because regulatory approval for release was contingent on limiting the introduction of alien genetic material and ensuring the continued insecticide susceptibility of Australian *Ae. aegypti* populations by excluding the possibility of transferral of insecticide resistance alleles.

## RESULTS

### *Wolbachia w*AlbB is widespread throughout mosquito but infection density varies between tissues

The density of *Wolbachia* within female *w*AlbB2-F4 mosquitoes was analysed by immunofluorescence analysis (IFA) using an antibody against the *Wolbachia* surface protein (WSP) and DAPI staining for DNA (Fig. 1). *Wolbachia* infection was widespread though internal mosquito tissues but the staining density (determined from the ratio of the area of *Wolbachia* WSP staining to mosquito cellular DNA) varied between tissues (Fig. 1F). Of several tissue types examined, *Wolbachia* staining density was highest in mosquito oocytes (Fig. 1B, F), followed by salivary glands (Fig. 1D, F). Interstitial spaces and heads had moderate staining densities (Fig. 1E and F). Significantly lower staining densities were observed in thoracic ganglia, flight muscles and midguts (Figs 1A, C and F).

**Figure 1.**
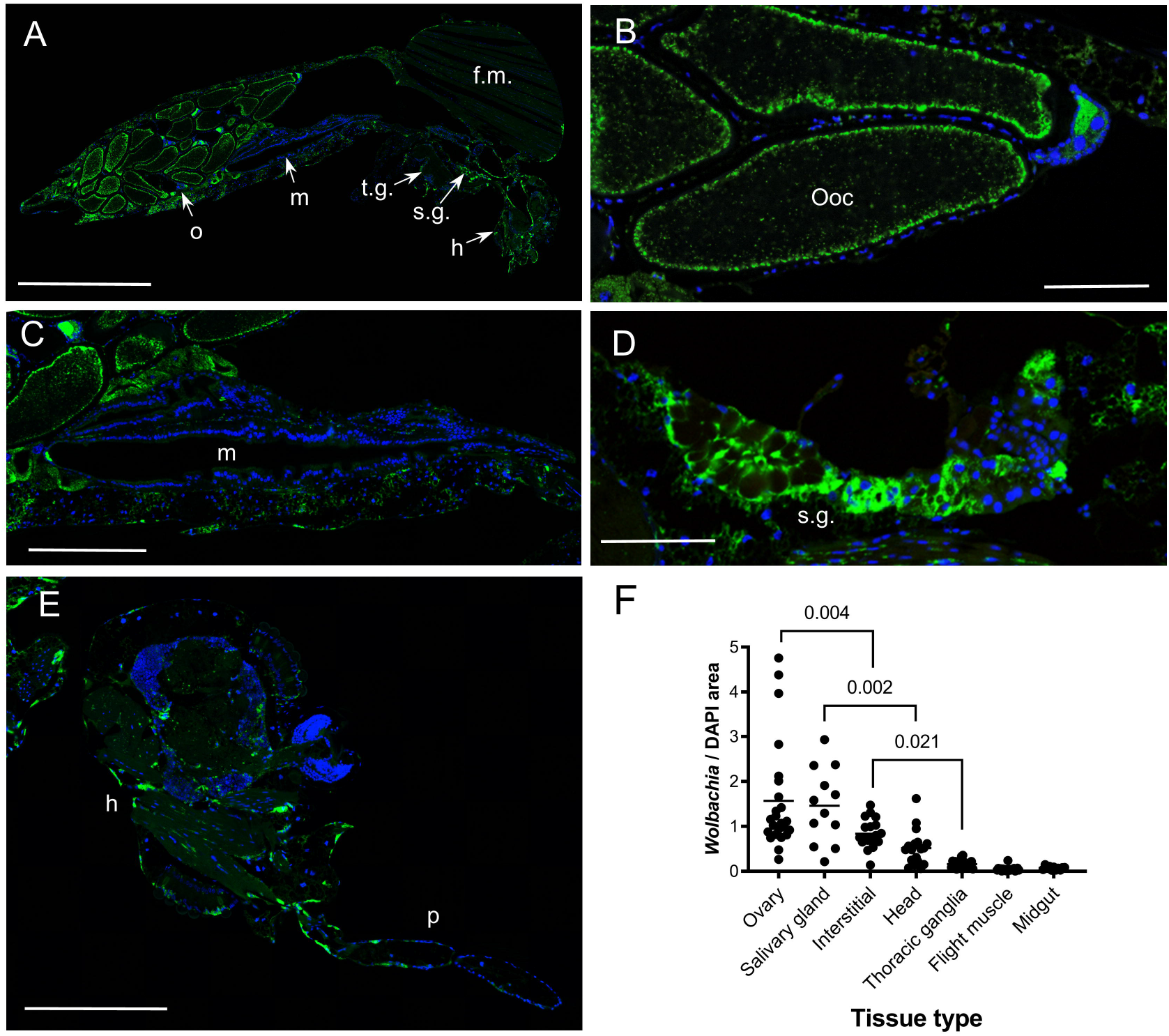
Histology of *Wolbachia* infection in the *Aedes aegypti w*AlbB2-F4 strain. *Wolbachia* infection was observed across mosquito organs/tissue types by immunofluorescence analysis (IFA) using a rabbit polyclonal antibody against the *Wolbachia* surface protein (WSP) as the primary antibody and Alexa Fluor 488-conjugated donkey anti-rabbit antibody as the secondary antibody. DNA was stained using DAPI. (A) Example of whole body section showing IFA staining. (B-E) High resolution images of *Wolbachia* staining in oocytes, midgut, salivary gland and heads, respectively. (F) Quantification of *Wolbachia* staining density. Staining areas were quantified by image analysis and expressed as a ratio of *Wolbachia* staining over DAPI staining for each organ/tissue. Green, *Wolbachia*, blue DNA. h, head. f.m., flight muscles. midgut. o, ovary. ooc, oocyte. p, proboscis. s.g. salivary glands. t.g., thoracic ganglia. Scale bars: A: 1.00 mm, B, D: 0.10 mm. C, E: 0.25 mm

### The *w*AlbB2-F4 strain is resistant to dengue and Zika viruses

We assessed the level of *Wolbachia-*induced suppression of DENV-2 and ZIKV in *w*AlbB2-F4 mosquitoes compared to Australian wild type (*Wolbachia*-free) *Ae. aegypti*. Both wild type and *w*AlbB2-F4 mosquitoes were fed a blood meal containing a strain of DENV-2 isolated from a dengue fever patient in Australia in 2015 (QML16) at a titre of 1 × 10^6.6^ CCID_50_/ml (in C6/36 cells) before being incubated at 28°C, 75% relative humidity and 12:12 hr day:night light cycle for 14 d post infection.

The presence and intensity of DENV infection in mosquito tissues was analysed by quantitative reverse transcriptase PCR (qRT-PCR) targeting a region of the DENV 3’ untranslated region (UTR) [36]. At 14 d post feeding (dpf), 69.2% of wild type mosquitoes were infected with DENV, whereas only 16.7% of *w*AlbB2-F4 mosquitoes were infected, representing a highly significant reduction due to *Wolbachia* infection (P < 0.0001, Fig 2A). Furthermore, there was a significant reduction in the percentage of mosquitoes with virus present within legs and wings (indicating a disseminated infection) in *w*AlbB2-F4 mosquitoes compared to wild type mosquitoes (P = 0.013, Fig. 2A). There was a highly significant reduction in the number of virus copies in the bodies (P < 0.001) and legs and wings (P = 0.0025) of *w*AlbB2-F4 mosquitoes compared to wild type mosquitoes (Fig. 2B). The presence and quantity of live virus in saliva was analysed by cell culture ELISA [40]. DENV was not detected in the saliva of *w*AlbB2-F4 mosquitoes, but it was detected in the saliva of 8.3% of wild type mosquitoes (Fig 2A). The relative infection density of *Wolbachia* and DENV was visualised using dual-antibody IFA of dengue and *Wolbachia* infection, using the *Wolbachia* WSP antibody, an antibody against flavivirus non-structural protein 1 (NS1) and DAPI staining. DENV infection could be observed in tissues throughout the head, thorax and abdomen of wild type mosquitoes (Fig 2C). In contrast, of the *w*AlbB2-F4 mosquitoes that were infected, virus was restricted to the midgut (Fig. 2D). Staining density was higher in the midguts of wild type mosquitoes (Fig 2E) compared with *w*AlbB2-F4 mosquitoes (Fig. 2F). DENV was observed in the salivary glands of wild type mosquitoes, however it could not be detected from salivary glands of *w*AlbB2-F4 mosquitoes (Fig. 2F). In *w*AlbB2-F4 females the restriction of DENV infection to the midgut corresponded with a low density of *Wolbachia* infection in that tissue (Figs. 2F and 1F).

**Figure 2.**
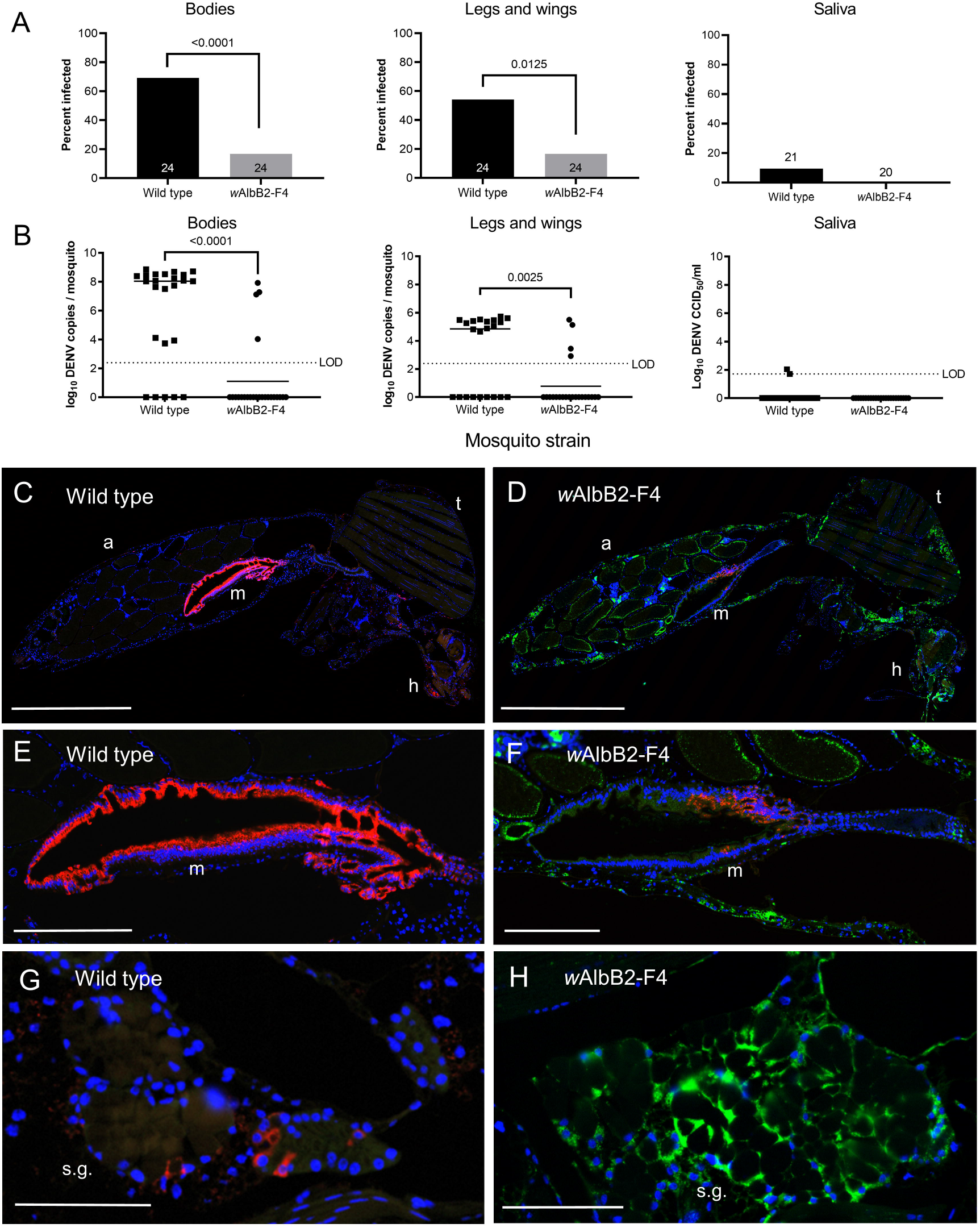
*Wolbachia* inhibit dengue 2 infections in *Ae. aegypti w*AlbB2-F4 mosquitoes. (A) DENV-2 virus prevalence in the bodies, legs and wings, and saliva of *Ae. aegypti w*AlbB2-F4 and wild type mosquitoes 14 d after feeding on a blood meal containing 1 × 10^6.6^ CCID_50_/ml (in C6/36 cells) of DENV-2 virus. (B) DENV-2 infection intensity in mosquitoes in A. Virus copy numbers were determined from bodies, legs and wings and saliva samples using quantitative reverse-transcriptase PCR (qRT-PCR). (C) Example whole body midsagittal section from a wild type mosquito dual stained for *Wolbachia* (green) and DENV-2 (red). (D) Whole body section of a *w*AlbB2-F4 female showing restriction of virus to the midgut. (E-F) High resolution images of midguts from wild type and *w*AlbB2-F4 mosquitoes showing lower DENV-2 staining density in the *w*AlbB2-F4 midgut. (G-H) High resolution images of salivary glands from wild type and *w*AlbB2-F4 mosquitoes showing dense *Wolbachia* infection and absence of DENV-2 infection in the latter. Scale bars: C-D, 1 mm. E-F, 0.25 mm, G-H: 0.10 mm.

Mosquitoes from the wild type and *w*AlbB2-F4 mosquitoes were also provided a blood meal containing a strain of Zika virus isolated from a febrile patient in Paraiba State during the 2015/2016 Brazil epidemic at a titre of 1 × 10^8.5^ CCID_50_/ml (in C6/36 cells) (Fig 3A). Mosquitoes were incubated for 14 dpi and the presence and intensity of Zika virus infection in mosquito segments was analysed by qRT-PCR [41]. All wild type and *w*AlbB2-F4 mosquitoes had detectable virus in bodies and legs and wings at 14 d post feeding (Fig. 3A). However, significantly fewer virus copy numbers were observed in both the bodies and legs and wing tissue of the *w*AlbB2-F4 mosquitoes at 14 dpi compared to wild type mosquitoes (P < 0.0001, Fig. 3B). Live ZIKV was detected and quantified in mosquito saliva using the NS1 antibody as described above. Importantly, there was a highly significant reduction (P = 0.005) in the proportion of *w*AlbB2-F4 mosquitoes that expectorated virus in saliva compare to wild type mosquitoes.

**Figure 3.**
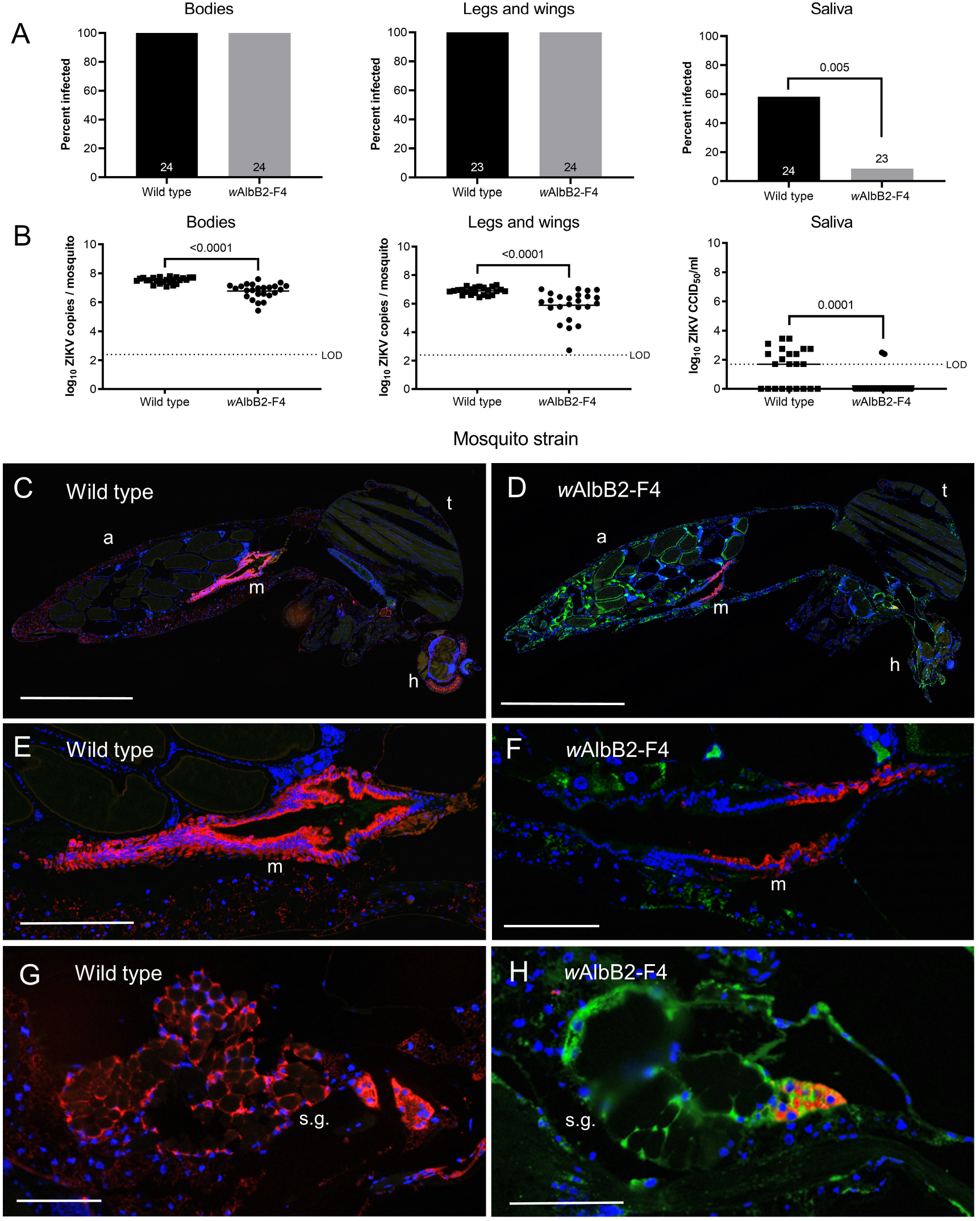
*Wolbachia* inhibit ZIKV infections in *Ae. aegypti w*AlbB2-F4 mosquitoes. (A) ZIKV virus infection prevalence in *Ae. aegypti* wild type and *w*AlbB2-F4 mosquitoes 14 d after feeding on a blood meal containing 1 × 10^8.5^ CCID_50_/ml (in C6/36 cells) of Zika virus. (B). ZIKV infection densities in mosquitoes from A. Virus copy numbers were determined from bodies, legs and wings and saliva samples using qRT-PCR. (C-D) Example whole body midsagittal sections of wild type and *w*AlbB2-F4 mosquitoes dual stained for *Wolbachia* (green) and ZIKV (red). (E-F) High resolution images of midguts from wild type and *w*AlbB2-F4 mosquitoes, respectively, showing relatively lower staining density for *w*AlbB2-F4 mosquitoes. (G-H) High resolution images of midguts from wild type and *w*AlbB2-F4 mosquitoes, respectively. Infection was limited and spatially restricted in *w*AlbB2-F4 mosquitoes. h, head. LOD, limit of detection. m, midgut. s.g. salivary gland. Scale bars: C-D, 1 mm. E-F, 0.25 mm, G-H: 0.10 mm.

Dual antibody IFA of ZIKV using the anti-WSP and Flavivirus NS1 antibody (above) revealed that, by 14 dpi, ZIKV had disseminated widely throughout mosquitoes from the wild type strain (Fig. 3C) but was generally restricted to midgut tissue in females from the *w*AlbB2-F4 strain (Fig. 3D). ZIKV infection in midgut tissue reached high densities in wild type females (Fig 3E) but was visibly lower in the midgut tissue of *w*AlbB2-F4 females. Similarly, the salivary glands of wild type females were observed to have dense ZIKV infection, but the infection was lower and spatially restricted in *w*AlbB2-F4 salivary glands (Fig. 3F). In *w*AlbB2-F4 females, the highest densities of ZIKV infection were observed in midgut tissue, in which the *Wolbachia* infection density was lowest (Figs. 3F and 1F).

### *w*AlbB2-F4 strain has a genomic background of the Australian *Aedes aegypti* and is susceptible to commonly-used insecticides

Double digest RAD genome-wide sequencing of mosquitoes from the *w*AlbB2-F4 strain and two strains used for its generation (WB2 strain and wild type Australian strain) revealed that the backcrossing procedure used resulted in *w*AlbB2-F4 mosquitoes having more than 98% wild-type Australian ancestry (Fig. 4A). Q values from the ADMIXTURE analysis show this ancestry percentage is consistent across all analysed *w*AlbB2-F4 individuals (Fig. 4A).

**Figure 4.**
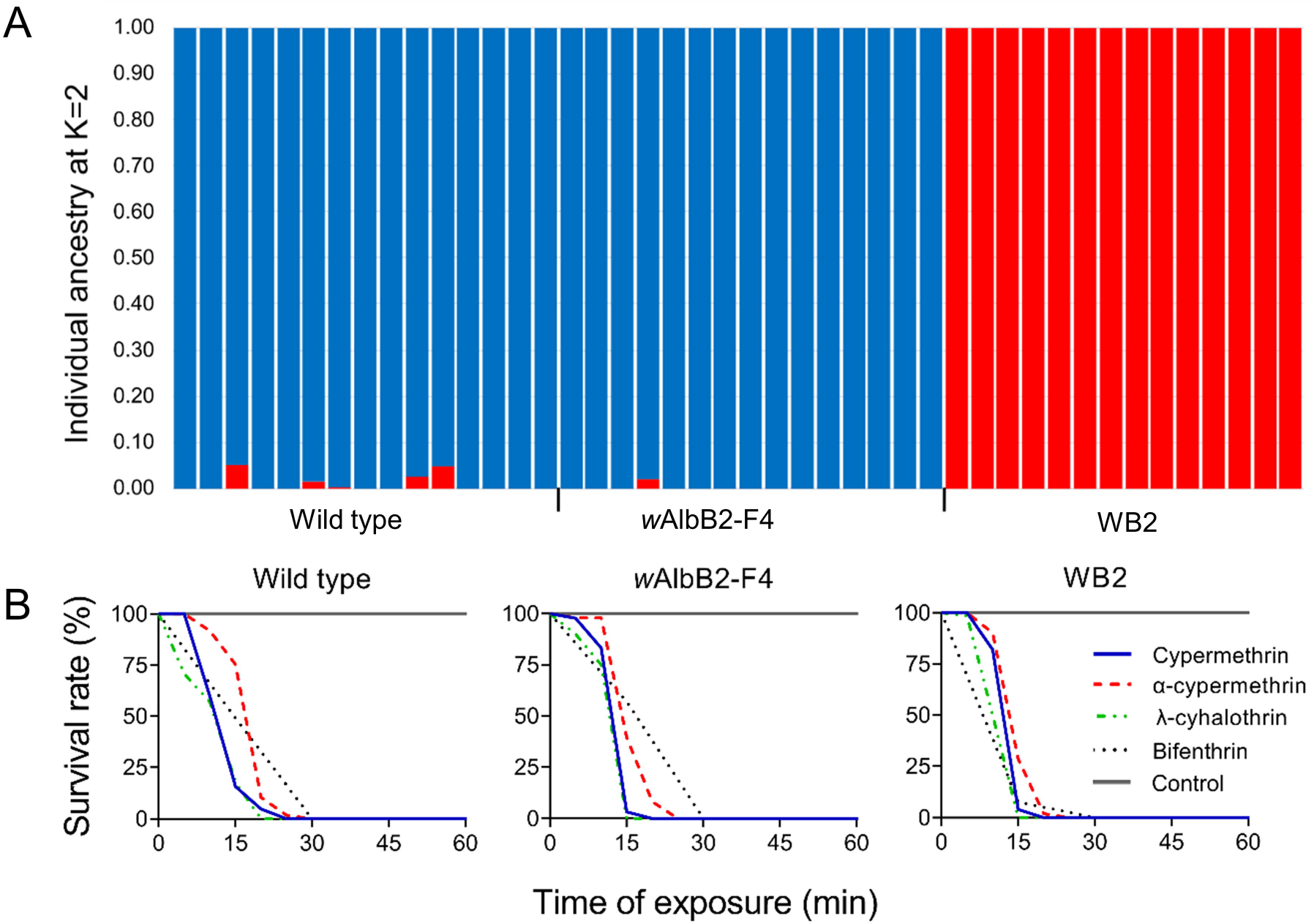
Analysis of genetic equivalency of *w*AlbB2-F4 strain to Australian *Aedes aegypti*. (A) All mosquitoes from the *w*AlbB2-F4 strain have >98% ancestry (Q-values) from the Australian wild-type strain (and not the WB2 strain), indicating a successful backcrossing procedure. (B) The *Wolbachia* infected *Aedes aegypti w*AlbB2-F4 strain has equivalent insecticide susceptibility to the parental Queensland wild type mosquitoes and *Ae. aegypti w*AlbB2 strains. Survival rates of mosquitoes were recorded following exposure to diagnostic doses of alpha-cypermethrin, cypermethrin and lambda-cyhalothrin using CDC bottle bioassays. Survival rates following exposure to a diagnostic dose of bifenthrin were determined using WHO filter paper assays.

Females from the *w*AlbB2-F4 strain and the parental wild type and WB2 strains exhibited rapid mortality from first exposure to the insecticides cypermethrin, α-cypermethrin, λ-cyhalothrin and bendiocarb (Fig. 4B). 100% (n = 40-82) of females died following treatment with the diagnostic doses of these insecticides by the diagnostic time of 30 mins, whereas the 30 min mortality of females in control tests (solvent only) was 0% for all tests. These results demonstrate that the *w*AlbB2-F4 and parental strains met the WHO definition of being susceptible to the insecticides tested. Insecticide resistance phenotypes were absent.

We also tested for the presence of insecticide resistance genotypes in these strains. The kdr mutations, V410L, V1016I and F1534C were not detected in the *w*AlbB2-F4 strain, nor the two parent strains, WB2 and Cairns WT, whereas the respective wild type alleles were amplified 100 % of successful qPCR reactions from individuals from these strains. In contrast, the kdr mutations were detected from >47% of mosquitoes from the *Ae. aegypti* Merida strain, which was consistent with the prevalence of kdr phenotypic expression in this strain (data not shown).

## DISCUSSION

We recently reported the generation of an Australian strain of *Ae. aegypti* infected with the *w*AlbB2 strain of *Wolbachia* that was used to successfully suppress a mosquito population in trial sites in northern Australia [15]. Here we show that the strain is also highly amenable for use in mosquito population replacement interventions; whereby persistent *Wolbachia* infections are established in target populations of mosquitoes to substantially reduce their capacity for transmitting pathogenic arboviruses [13, 24]. Compared to wild type (*Wolbachia*-free) mosquitoes, 76% fewer *Ae. aegypti* from this (*w*AlbB2-F4) strain became infected following a blood meal that contained a contemporary DENV-2 virus stock. Significantly fewer *w*AlbB2-F4 mosquitoes developed a disseminated infection and virus could not be detected from saliva expectorates. Females from the *w*AlbB2-4 and wild type strains were susceptible to infection from a high dose of Zika virus, however, 85% fewer *w*AlbB-2 produced virus in saliva (a proxy for transmission potential) compared to the wild type strain. We showed that the *w*AlbB infection follows a typical pattern through different tissues, including high a high density in oocytes, and that high flavivirus infection density in midgut tissue is associated with low tissue density of *Wolbachia* infection. We also showed that the *Ae. aegypti w*AlbB2-F4 strain has an equivalent genetic background to wild mosquitoes from Queensland, Australia, having >98% Queensland ancestry as estimated from genome-wide SNP markers, and that susceptibility to common insecticides present in wild-type Australian mosquitoes is maintained in our *w*AlbB2-F4 strain. Equivalency to target populations of mosquitoes in genotype and insecticide susceptibility is critical for the successful invasion of *Wolbachia* into wild mosquito populations for population replacement [42] and can be a prerequisite for regulatory approval in jurisdictions with strong biosecurity controls, such as Australia.

There was a four-fold reduction in the proportion of *Ae. aegypti w*AlbB2-F4 mosquitoes that became infected with a contemporary DENV-2 strain (16.7%) compared to wild type mosquitoes (69.2%) and a three-fold reduction in the proportion of mosquitoes that developed a disseminated infection (16.6 % in *w*Ablb2-F4 against 54.2% in wild-type). DENV-2 loads in bodies and legs and wings were highly significantly reduced by *Wolbachia* infection. The reduced susceptibility of *w*AlbB2-F4 mosquitoes to infection and dissemination of DENV correlates well with results from an independently-generated *Ae. aegypti* strain transinfected with *w*AlbB *Wolbachia* (referred to as the *Ae. aegypti* WB1 strain) [37] and for *w*AlbB introgressed into a Taiwanese *Ae. aegypti* background in which all four DENV serotypes were inhibited [43]. We were also able to reveal differences in tissue distribution between *Wolbachia* and DENV-2 infection by performing dual antibody immunofluorescence analysis. Infections in wild type mosquitoes at 14 d post feeding were characterized by intense staining in midguts and isolated pockets of disseminated infection throughout the body. Conversely, in *w*AlbB2-F4, DENV-2 infection was restricted to midgut tissue, which had the lowest tissue densities of *Wolbachia* infection, in keeping with reported positive relationship between *Wolbachia* density and virus inhibition for most *Wolbachia* strains [44]. As the mosquito midgut is the first tissue to encounter arbovirus from an infected blood meal, the establishment of virus infection in these mosquitoes may be been due to the absence of *Wolbachia* in midgut tissue, whereas the inhibition of virus dissemination may have been due to the widespread *Wolbachia* infection in surrounding tissues.

For a high dose of an epidemic strain of Zika virus (2015 epidemic in Brazil) all mosquitoes from *w*AlbB-F4 and wild type strains developed primary and disseminated infections. However, there was a highly significant reduction in the proportion of *w*AlbB2-F4 mosquitoes producing virus in saliva (8.7% compared to 58.3% for wild type). The infection of all tested mosquitoes was likely due to the very high titre of Zika virus fed to mosquitoes (10^8.5^ CCID_50_/ml [in C6/36 cells]). The antiviral effect of *Wolbachia* infection was evident from the significantly lower ZIKV loads in bodies and legs and wings of *w*AlbB2-F4 mosquitoes, and the substantially lower proportion of *w*AlbB2-F4 mosquitoes that produced detectable ZIKV in saliva. The presence of virus in saliva is a proxy for the ability of a mosquito to transmit the virus and is therefore the most epidemiologically relevant measure. Our results correlate closely to results from similar experiments in other *Ae. aegypti* transinfected with the *Wolbachia w*AlbA [45] and *w*AlbB [14] strains. For both strains, *Wobachia* infection was associated with modest reductions to mosquito infection and dissemination rates but induced complete blockage of virus transmission (no virus was detected from salivas). Their results were also supported by our dual immunofluorescence analysis of ZIKV and *Wolbachia*. DENV and ZIKV was generally restricted to midgut tissue in the *w*AlbB2-F4 strain.

We confirmed that the *w*AlbB2-F4 strain is genetically equivalent to wild type *Ae. aegypti* from Queensland. For the maximally-divergent parental strains, the backcross mating strategy used for the creation of *w*AlbB2-F4 should ideally transfer >90% of the Australian genome [46] and we achieved an actual rate of transfer of 98%. Matching the genetic background of released and target mosquitoes is critical for the success of mosquito population replacement interventions [47]. It increases the likelihood that the released mosquitoes will survive and mate competitively within the target population, providing the best chance for persistence of *Wolbachia* infection and therefore the virus blocking phenotype. It is particularly important that released mosquitoes have equivalent insecticide susceptibility to the target population. Insecticides remain the primary means of mosquito control and various genetically determined mechanisms of insecticide resistance have evolved within *Ae. aegypti* populations. Releases of *w*MelBr-infected *Ae. aegypti* failed to achieve population replacement in Rio de Janeiro due to an absence of insecticide resistance in this strain in an environment where there is high household use of insecticides [42]. These conditions had selected for resistance in the resident population. Once the infected strain was backcrossed with resistant mosquitoes from Rio de Janeiro (*w*MelRio), the replacement intervention was successful (27). Genetic equivalency between the released and target mosquitoes is also important in particular circumstances where the jurisdiction intended for implementation of *Wolbachia* releases has strict regulations prohibiting the release of exotic organisms, which is the case in Australia and other isolated nations in Oceania. We have provided a new example that that introgression of *w*AlbB2 infection can be achieved by a straightforward process of back cross mating that transfers the *Wolbachia-*induced virus blocking phenotype into a mosquito background that is equivalent to the target population [43].

A pertinent question is whether *Wolbachia* strains other than the widely distributed *w*Mel strain are necessary for future population replacement interventions. While *Wolbachia w*Mel infections have persisted in *Ae. aegypti* at release sites in northern Queensland over a period of nearly a decade, recent laboratory and field evidence indicates that the *w*Mel infections are susceptible to heat stress [32-34]. The persistence of *w*Mel infections at release locations for over a decade has been attributed to the natural behaviour of mosquitoes to rest in locations that provide suitable microhabitats [26]. However, if the current trend of increasing global temperatures due to climate change continues [48], the availability of suitable microhabitats may be depleted.

The *Wolbachia w*AlbB strain has great potential to be applied in mosquito population replacement interventions against flavivirus diseases based on traits demonstrated here and elsewhere, including; induction of substantial reductions to flavivirus vector competence of transinfected mosquitoes [45, 49], induction of complete maternal inheritance and cytoplasmic incompatibility [15], ease of introgression into target mosquito genetic backgrounds and tolerance of heat stress [14, 33].

## METHODS AND MATERIALS

### Mosquito strains

Wild type *Ae. aegypti* were established from egg collections made in Cairns and Innisfail in 2015 and 2016, respectively, and they were confirmed to be uninfected with *Wolbachia* by PCR (data not shown). Establishment of the *Ae. aegypti w*AlbB2-F4 strain was described in [15]. Briefly, matings were set up with one male from the wild type Australian strain and three virgin females from the USA WB2 strain, imported from the Michigan State University. These females were then blood fed and allowed to lay eggs. The F1 eggs were hatched, female pupae separated and reared till adulthood to be then mated with wild type males. The procedure was repeated for two additional generations to obtain the F4 generation of the backcross refer to as the *w*AlbB2-F4 strain. The mosquito colonies were maintained in the QIMR Berghofer insectary at 28°C, 70% relative humidity and 12:12 hr light cycling with dawn and dusk fading. Adults were maintained in 30 × 30 × 30 cm cages (BugDorm, MegaView Science Education Services Co., Ltd., Taiwan) and provided with 10% sucrose solution *ad libitum* and defibrinated sheep blood (Serum Australis, Manila, NSW, Australia) once per week. To provide mosquitoes for experiments, eggs were flooded in plastic trays and larvae were maintained at a density of 500 larvae in 4 L of aged tap water and fed ground TetraMin tropical fish food flakes *ad libitum* daily before the resulting pupae were sorted into adult emergence trays and transferred to cages.

### Viruses

A strain of DENV-2 (QML16) originally isolated from a dengue fever patient in Australia in 2015 was provided by Prof John Aaskov, Queensland University of Technology, Australia. A strain of ZIKV (KU365780) isolated from a Zika virus disease patient in Joao Pessoa, Paraiba State, Brazil on 18 May 2015 was provided by Pedro Fernando da Costa Vasconcelos, Evandro Chagas Institute, Brazil. Both viruses were propagated in C6/36 cells at 28°C, 5 % CO_2_ for 5 d. Infected cell culture supernatants were harvested for these experiments. ZIKV was concentrated using an Amicon Ultra-15 Centrifugal Filter Unit with an Ultracel-100 membrane (Merck Millipore, Darmstadt, Germany).

### Vector competence of the *Ae. aegypti* wAlbB2-F4 strain for dengue and Zika viruses

#### Mosquito infection

Approximately 100 *w*AlbB2-F4 or wild type females were placed in 750 ml plastic containers with gauze lids and fed mixtures of DENV-2 or Zika virus in defibrinated sheep blood (Serum Australis) using glass artificial membrane feeders [50]. Blood virus mixtures consisted of DENV-2 supernatant and defibrinated sheep blood at a ratio of 1:1, or ZIKV stock and defibrinated sheep blood at a ratio of 1:5 and samples of the blood virus mixtures were taken before and after the feeding period to determine virus titres. After the feeding opportunity, all mosquitoes were anaesthetised with CO_2_ and placed on a petri dish on ice. Non- or partially engorged mosquitoes were discarded and fully engorged mosquitoes were placed in containers, provided with 10% sugar solution *ad libitum* and housed in an environmental chamber (Panasonic, Osaka, Japan) set at 28°C, 75% RH, and lighting conditions described above. Mosquitoes were harvested 14 d after blood feeding, anaesthetized and placed on ice. Legs and wings were removed and placed into 2 ml screw cap vials with three 2.3 mm zirconium silica glass beads. Saliva was collected by placing mosquito bodies on double sided tape and positioning a 200 ml pipette tip containing 10 µl of saliva collection fluid (10% FBA, 10% sugar [51]) over the proboscis of each mosquito for 20 min. The contents were expelled into a 1.5 ml microfuge tube. Each body was placed into a 2 ml screw-cap tube with beads as described above.

#### Quantification of the virus from mosquito bodies

Virus nucleic acid was extracted using the Roche High Pure virus nucleic acid extraction kit by adding 200 µl of the working binding buffer to the tubes containing bodies or legs & wings and homogenizing the tissues by shaking the tubes for 1 min 30 s using a Mini Beadbeater-96 (BioSpec Products, Bartlesville, OK, USA). The tubes were centrifuged for 8,000 × g for 1 min. 50 µl Proteinase K was added and the procedure was continued as described in the manufacturer’s protocol.

Dengue virus quantification was performed by One-step RT-qPCR using the Taqman Fast Virus 1-Step Master Mix and primers and probe targeting the DENV 3’UTR region described by Frentiu et al [36]. Ten µl reactions contained 2.5 µl of 4 x Taqman Fast virus mastermix, 400 nM of each primer, 250 nM of probe and 1 µl of virus nucleic acid extraction. Primers and probe were synthesized by Macrogen (Macrogen, Seoul, Korea). Thermal cycling was performed using a Corbett Rotorgene 6000 (QIAGEN/Corbett, Sydney, NSW, Australia) with incubation at 50°C for 5 min, 95°C for 20 s, then 40 cycles of 95°C for 3 s and 60°C for 30 s. Absolute quantification of virus copy number was performed using the Rotorgene 6000 software package using a standard curve derived from 10-fold serial dilutions of a linearized plasmid containing the 3’UTR gene [36]. Zika virus quantification was performed by One-step RT-qPCR using the reaction mix and thermocycling conditions described above but with primers ZIKV 911C and ZIKV 835 and probe ZIKV 860-FAM [41]. A 10-fold serial dilution series of a linearized plasmid containing the target sequence [40] was used for determination of Zika virus copy number.

#### Cell culture ELISA for determination of live virus in blood meals and mosquito saliva

Blood virus mixtures were titrated by ten-fold serial dilution in virus media (RPMI 1640 cell culture media supplemented with 5% fetal bovine serum [FBS] and 1% Penicillin-Streptomycin [Gibco]) in a 96-well plate before transferring dilutions to equivalent wells of a 96 well plate containing near confluent C6/36 cell monolayers. The inoculated cells were incubated at 28°C, 5% CO_2_, for five d. Five-fold dilutions of saliva samples were tested and the inoculated cells were incubated for six d. After incubation, cell monolayers were fixed by adding 100 µl of ice cold 80% acetone / PBS and incubating plates at -20°C for 1 hr. The acetone was removed and plates were rinsed three times in PBS. DENV and Zika antigen was detected by performing an Enzyme Linked Immunosorbant Assay (ELISA) targeting *Flavivirus* NS1 protein. Fixed cells were blocked in 100 µl of blocking buffer (1% [w/v] bovine serum albumin in PBS) at room temperature for 1 hr. Cells were then washed three times in PBS/0.05% Tween 20 (PBS-Tween). The cells were incubated with 50 µl 4G4 anti-Flavivirus NS1 monoclonal hybridoma supernatant[52] (1:40 in PBS-Tween) and then washed three times in PBS-Tween. Cells were incubated with 50 µl Horse Radish Peroxidase (HRP-) conjugated goat anti-mouse antibody (Dako) (1:2000 in PBS-Tween) before being washed four times in PBS-Tween 20. Plates were dried and wells were incubated with 50 µl of Tetramethylbenzidine (TMB) Liquid Substrate for Membranes (Sigma Aldrich) for 30 min at room temperature. Blue staining of the cell monolayer indicated the presence of virus infection in cells. The 50% Cell Infectious Dose (CCID_50_) per ml in C6/36 cells was calculated using the method of Reed and Muench [53].

#### Histological analysis of Wolbachia and flavivirus infections

Infections of *Wolbachia* and DENV-2 or ZIKV were detected in thin paraffin sections from mosquitoes by immunofluorescence analysis based on established protocols [40] with the following modifications for dual staining of *Wolbachia* and flaviviruses. Nonspecific antibody binding by incubating mosquito sections in 10% donkey serum for 60 min. Excess serum was decanted and the first primary antibody, 4G4 mouse anti-Flavivirus NS1 [52], was applied undiluted overnight in a humidified chamber. Sections were washed three times in Tris buffered saline plus 0.025% Tween 20 (TBS_TW_).10% donkey serum was applied for 15 min before a rabbit anti-WSP polyclonal antibody diluted 1:500 [44] in 10% donkey serum was applied for 2 h at room temperature in a humidified chamber. Sections were washed three times in TBS_TW_. Alexa Fluor donkey anti-mouse 555 diluted 1:300 and Alexa Fluor 488 donkey anti-rabbit diluted 1:1000 in TBS was applied for 60 min. Sections were washed three times in TBSTw. Sections were counterstained with DAPI for 10 min, washed several times in PBS and mounted with Vector Vectashield or Dako Fluorescence Mount. Microscopy was performed using an Aperio ScanScope fluorescent microscope using filters for DAPI, Alexa 555 (Cy 3) and Alexa 488 (FITC) and exposure times of 0.1 s, 0.2 s and 0.16 s, respectively. The relative staining density of *Wolbachia* to DAPI stained DNA was measured digitally from images using established protocols [40].

### Genome-wide characterization of the *w*AlbB2-F4 and parental strains

#### DNA extraction, sequencing, and genotyping of individual mosquitoes

Total genomic DNA was extracted from individual mosquitoes using DNeasy Blood and Tissue DNA extraction kit (Qiagen, Hilden, Germany) according to the manufacturer’s instructions. We prepared one double-digest RADseq library with DNA from 44 individually-barcoded mosquitoes: 15 from the *w*AlbB2-F4 strain, 15 from the wild-type Australian strain (Cairns), and 14 mosquitoes from the WB2 strain. The library was prepared following the protocol described in Rašić et al. (https://doi.org/10.1186/1471-2164-15-275), and sequenced on one lane of the Illumina HiSeq4000 platform. The sequencing data were demultiplexed and processed (trimmed to 90□bp and filtered for quality) using the bash script/pipeline from Rašić et al. (https://doi.org/10.1186/1471-2164-15-275). The high-quality reads were aligned to the *Ae. aegypti* genome assembly version AaegL5 (https://doi.org/10.1038/s41586-018-0692-z) with the aligner Bowtie (https://doi.org/10.1186/gb-2009-10-3-r25). Unambiguously mapped reads were converted to the bam format and processed in SAMtools (DOI: 10.1093/bioinformatics/btp352). The sorted bam files were passed to the ANGSD pipeline, where the SAMtools algorithm was used for variant and genotype calling (https://doi.org/10.1186/s12859-014-0356-4). The final VCF file contained 12931 SNP markers that were present in at least 75% of individuals, had a minor allele detected in at least 2 individuals, and gave genotype likelihoods of at least 95%.

#### ADMIXTURE analysis

To assess if the backcrossing procedure resulted in the expected high genome-wide similarity between the *w*AlbB2-F4 strain and the Australian wild type strain, we performed ADMIXTURE analysis that estimates ancestry proportions for each individual (DOI: 10.1101/gr.094052.109). To avoid estimation bias caused by the highly-linked markers, we pruned SNPs so that they are at least 100 kb apart using VCFtools (doi: 10.1093/bioinformatics/btr330), and ran ADMIXTURE analysis with 3803 unlinked SNP markers (distributed across all three *Ae. aegypti* chromosomes) while assuming two ancestral populations (K=2).

### Insecticide resistance bioassays

Each of the three *Ae. aegypti* strains (wild type Australia, WB2 and *w*AlbB2-F4) was assessed for insecticide resistance to cypermethrin, alpha-cypermethrin and lambda-cyhalothrin and biphenthrin, the active constituents of commercially available insecticides in Queensland, Australia. Cypermethrin, alpha-cypermethrin and lambda-cyhalothrin were tested using CDC bottle bioassays. These were performed in insecticide coated glass bottles at their diagnostic dose using acetone as a solvent as per CDC guidelines (https://www.cdc.gov/malaria/resources/pdf/fsp/ir_manual/ir_cdc_bioassay_en.pdf). Females of each strain were divided and allocated into 4-7 treatment bottles (insecticide coated) and 1-2 control bottles (acetone only). Knock down was recorded every five min. After 120 min, mosquitoes were transferred to untreated containers, provided with 10% sucrose ad libitum, and maintained for 24 hr to assess recovery. Susceptibility to bifenthrin was tested using the WHO filter paper assay (https://apps.who.int/iris/bitstream/handle/10665/250677/9789241511575-eng.pdf) at the diagnostic dose of 0.025% and mortality scored at 15 min intervals.

Mosquitoes from Cairns WT, WB2, *w*AlbB2-F4 strains were tested for the presence of mutations in the voltage-gated sodium channel protein (VGSC) gene associated with knock down resistance to pyrethroids using Allele-specific quantitative PCR and melting curve analysis (AS-PCR) [54, 55]. These mutations included single nucleotide mutations causing changes from valine (V) to leucine (L) amino acid substitution at locus 410 (V410L), V to isoleucine (I) at locus 1016 (V1016I) and from phenylalanine (F) to cysteine (C) at position 1534 (F1534C) (numbered according to the homologous locus in the VCSC gene in *Musca domestica*). A strain of *Ae. aegypti* from Merida, Mexico, with partial expression of kdr resistance phenotypes was used as a positive assay control.

## ACKNOWLEDGEMENTS

Australian National Health and Medical Research Council (NHMRC 1082127) Staff at the Department of Agriculture and Water Resources and the Australian Pesticide and Veterinary Medicines Authority.

We thank Pedro Fernando da Costa Vasconcelos, Evandro Chagas Institute, Brazil, for provision of Zika virus and John Aaskov, Queensland University of Technology, Australia, for dengue virus stocks. We thank Roy Hall and Jody Hobson-Peters for the provision of anti-flavivirus antibody.

## AUTHOR CONTRIBUTIONS

## DECLARATION OF INTEREST

The authors declare no competing interest

## Notes

### Competing Interest Statement

The authors have declared no competing interest.

